# Alamar Blue assay optimization to minimize drug interference and inter-assay viability

**DOI:** 10.1101/2023.03.16.532999

**Authors:** Mina N. Dinh, Masahiro Hitomi, Zahraa A. Al-Turaihi, Jacob G. Scott

## Abstract

Alamar Blue (AB) has become an increasingly popular reagent of choice for cell viability assays. We chose AB over other reagents such as MTT and Cell-Titer Glo due to its cost effectiveness and its ability to be a nondestructive assay. While analyzing the effect of osimertinib, an EGFR inhibitor on the non-small cell lung cancer cell line, PC-9, we noticed unexpected right-shifts of dose response curves as compared to the curve obtained by Cell Titer Glo assay. Here, we describe our modified AB assay method to avoid right shift right shift in dose response curve. Unlike some of the redox drugs that were reported to directly affected AB reading, osimertinib itself did not directly increase AB reading. Yet, the removal of the drug containing medium prior to AB addition eliminated falsely increased reading giving comparable dose response curve as the one determined by Cell Titer Glo assay. When a panel of 11 drugs were assessed, we found that this modified AB assay eliminated unexpected similar right shifts detected in other epidermal growth factor receptor (EGFR) inhibitors. We also found that plate-to-plate variability can be minimized by adding an appropriate concentration of rhodamine B solution to the assay plates to calibrate fluorimeter sensitivity. This calibration method also enables a continuous longitudinal assay to monitor cell growth or recovery from drug toxicity over time. Our new modified AB assay is expected to provide accurate *in vitro* measurement of EGFR targeted therapies.

## Introduction

The Alamar Blue (AB) cell viability assay harnesses the optical property change associated with the conversion of resazurin (oxidized, non-fluorescent, blue) to resorufin (reduced, fluorescent, pink) that takes place under the cellular reducing environment. Resazurin is water-soluble, stable in culture medium, non-toxic and permeable through the cell membrane, making it optimal for measuring cell viability^1^. Resazurin can be reduced by NADH and NADPH in the presence of NADPH dehydrogenase or NADH dehydrogenase enzyme, and this reaction is thought to occur in the cytoplasm. Analysis of drug effect by this assay, however, may require caution because Alamar Blue has been reported with several interferences, including cell-culture media^2^ and anti-oxidant drugs^3^. Studies have suggested modifications that allow Alamar Blue or resazurin-based cell viability assays to be better suited and more reproducible, such as in 2D cell cultures^4^, 3D spheroid cultures^5^, and microbial biofilm staining^6^.

Osimertinib is a 3rd generation EGFR inhibitor and the newest FDA approved adjuvant therapy for non-small cell lung cancer (NSCLC) with EGFR exon 19 deletions or L858R mutation in exon 21. Acquired resistance to early generations of EGFR tyrosine kinase inhibitors (TKIs) is mostly driven by a second-site EGFR T790M mutation, which negates their inhibitory activity^7^. Osimertinib targets the T790M mutant EGFR by binding irreversibly to the C797 amino acid forming a covalent bond while sparing wild-type EGFR. Therefore, accurate measurement of Osimertinib sensitivity is critical to understand evolutionary dynamics of therapeutic resistance in NSCLC.

The cytotoxic effect of osimertinib on PC-9 cells was analyzed using Alamar Blue assay to understand the evolution of resistance in NSCLC against this EGFR inhibitor. We chose Alamar Blue based on its cost effectiveness and its potential as a non-destructive assay. Despite no report on Alamar Blue assay interference by osimertinib, we noticed an unexpected right shifted dose response curve resulting in a higher IC50 value than that determined by CellTiter-Glo (CTG), a luciferease based assay. This paper describes a simple method to eliminate this drug interference when using the Alamar Blue assay and to minimize inter-plate variability. We also demonstrate the utility of the optimized method to monitor the recovery from transient exposure to osimertinib.

## Results

### Drug presence interferes with the Alamar Blue assay

Osimertinib’s effect on PC-9, a non-small cell lung cancer cell line, was analyzed by cell viability assay using Cell Titer Glo (CTG), a luciferase based assay, and Alamar Blue. While optimizing AB assay incubation time, we noticed unexpected right shifts in the dose response curves (**Figure 1A)**. We cannot reliably assess drug resistance evolution if IC50 varies depending on the assay conditions. The degree of the shift was reduced by shortening incubation time, yet the IC50 value determined by the CTG assay was significantly lower than determined by the AB assay with 2 hr incubation, the shortest duration we tested (**Figure S1A & B)**.

**Figure 1.**
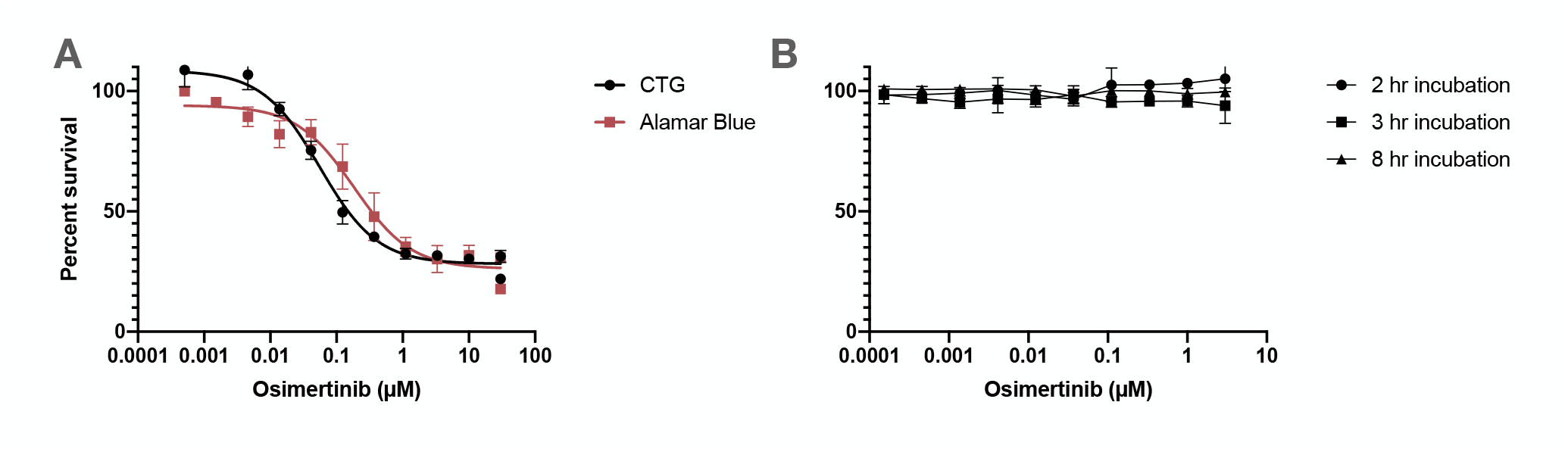
Osimertinib addition changes the Alamar Blue reading, but is not a direct cause. **(a)** Osimertinib increases the Alamar Blue reading when compared to Cell-Titer Glo. **(b)** Osimertinib did not directly increase Alamar Blue readings. PC-9 cells were plated into a 96 well plate at the density of 12,000 cells / 80 µl / well. After overnight culture, the cells received varying concentrations of osimertinib in 10 µl medium, and one tenth volume of Alamar Blue solution and incubated for 2, 3, and 8 hours. The Alamar Blue conversion reaction was stopped by adding 50 µl of 3% (w/v) SDS. The fluorescence intensity of each well was determined by fluorometry using BioTek Synergy H1 plate reader with excitation at 560 nm and emission at 590 nm. Mean + SD, n=3.

The AB assay relies on redox dependent conversion of resazurin to resorufin, and some drugs altering cellular redox state have been reported to interfere with AB assay^3^. In order to test if osimertinib has similar direct effect on the AB assay, a constant number of PC-9 cells were incubated with various concentrations of osimertinib together with AB for 2 hours. Unlike the data obtained after 5 days of osimertinib incubation, the presence of osimertinib only with AB did not interfere with florescence readings over 2, 3, and 8 hours**(Figure 1B)**. These results indicate that the AB assay interference by osimertinib is not a direct effect of the drug itself, and the possibility of AB assay interference cannot simply be predicted by direct drug addition to the assay.

The shift of response curve in AB assay resulted in a higher IC50 value when compared to those from CTG assay even when the incubation time was shortened to 2 hrs, the shortest duration we tested and the most similar to CTG (**Figure 2A**). The exact mechanism how osimertinib interferes AB assay is not clear, but removal of 5 day-drug treatment medium and addition of new medium with AB eliminated right shift of dose response curve. In contrast to AB assay, the dose response curve, hence IC50, determined by CTG was not affected by prior removal of drug treatment medium. We also observed various degree of right shifts of dose-response curves of other EGFRis (gefitinib, erlotinib and lapatinib) when AB assay was performed without removal of the drugs (**Figure 2A**). In all cases, the dose-response curves determined by AB assay after removal of drug treatment medium were comparable to the CTG assays, and the calculated IC50s are listed in **Figure 2B** with a depiction of the IC50s scaled to CTG values **Figure 2C**. Futhermore, we treated PC-9 cells with the IC50 calculated from the assay in **Figure 2D** and showed there was a statistically significant difference between AB with the presence of drug vs AB without. Removing drug treatment medium prior to assay, may preemptively eliminate unpredictable drug interference on AB assay. To eliminate drug interference, we found discarding the treatment medium is enough, and no washing was required (**Supplementary Figure S1**).

**Figure 2.**
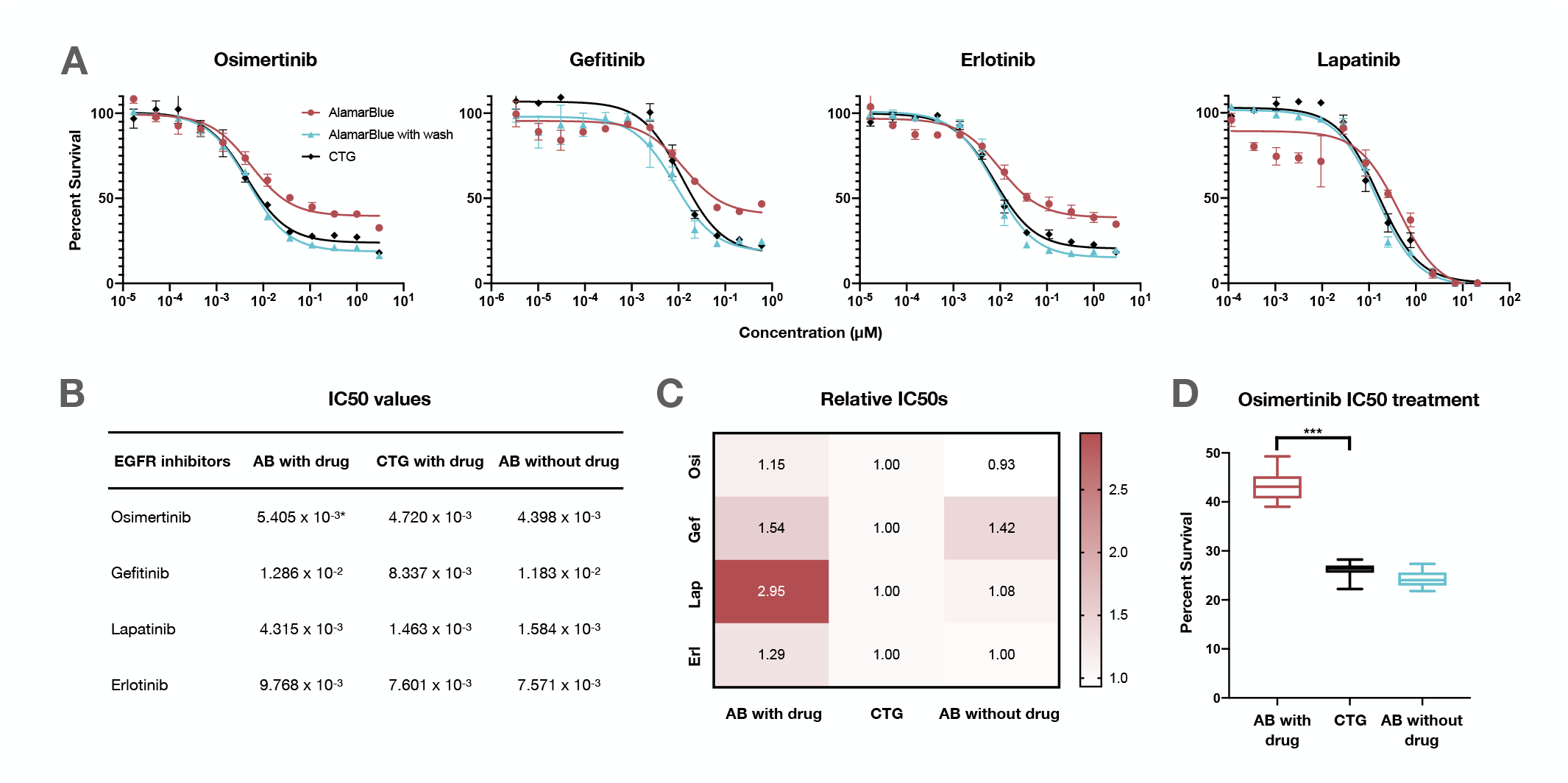
Alamar Blue assay interference by four EGFR inhibitors. **(a)** Four EGFR inhibitors show a shift that is resolved with “washing” the Alamar Blue media. **(b)** Listed IC50 concentrations calculated from panel (a). **(c)** Heatmap showing relative IC50s scaled to the CTG condition from panel (b). **(d)** IC50 values determined by AB assay in the presence of osimertinib was higher than those determined by CTG in the presence of osimertinib or by AB assay after removal of osimertinib. ***p=1.44e-12, n=21.

Interestingly, we found that dumping the media in our CTG condition also changed the expected drug curve, albeit a left shift. CTG is a luciferase assay that measures ATP content. We hypothesized that the lower reading was caused by the loss of ATP released into the medium when discarding the culture medium. To test this hypothesis, we separately measured the luminescence values of the medium to be discarded and that of corresponding wells after medium removal. When we added those two values together for each individual well, the curve based on the sum showed the same curve as CTG as a direct addition as as our hypothesis predicted (**Supplementary Figure S2**). This observation indicate that the presence of osimertinib does not interfere CTG assay.

### SDS quenches reduction of resazurin to resorufin

The stability of assay signal can be an important factor to perform multi-well plate based assays, as the availability of plate reader is often limited in the real research laboratory setting. We tested the stability of the AB fluorescence signal by reading the drug assay plates kept in dark at room temperature following 2 hour AB reaction in the absence of drug; using two drugs, osimertinib and gefitinib, another EGFR-TKI approved for EGFRmut lung cancer. The fluorescence intensity increased over time for assay plates of both drugs (**Figure 3A & C**). This time dependent increase might be due to the continuous conversion of resazurin to resorufin by the cells after the plates were taken out of the incubator. Indeed, this time dependent increase was quenched by addition of SDS (final concentration 1%) to wells to stop AB conversion reaction by lysing the cells, an optional step from the protocol published by Invitrogen (https://assets.thermofisher.com/TFS-Assets/LSG/manuals/AlamarBluePIS.pdf). We still detected a slight time-dependent increase of fluorescence signal sifting drug-response curves even after addition of SDS, but the calculated IC50s were not significantly different between each time step (**Figure 3B & D**). These data indicate that SDS addition should be be included to the protocol to estimate IC50 values as accurately as possible. Particularly when multiple plates are to be read, SDS addition would minimize plate-to-plate variation by instantly quenching the cellular activity that otherwise gradually increases the reading by converting resazurin to resorufin.

**Figure 3.**
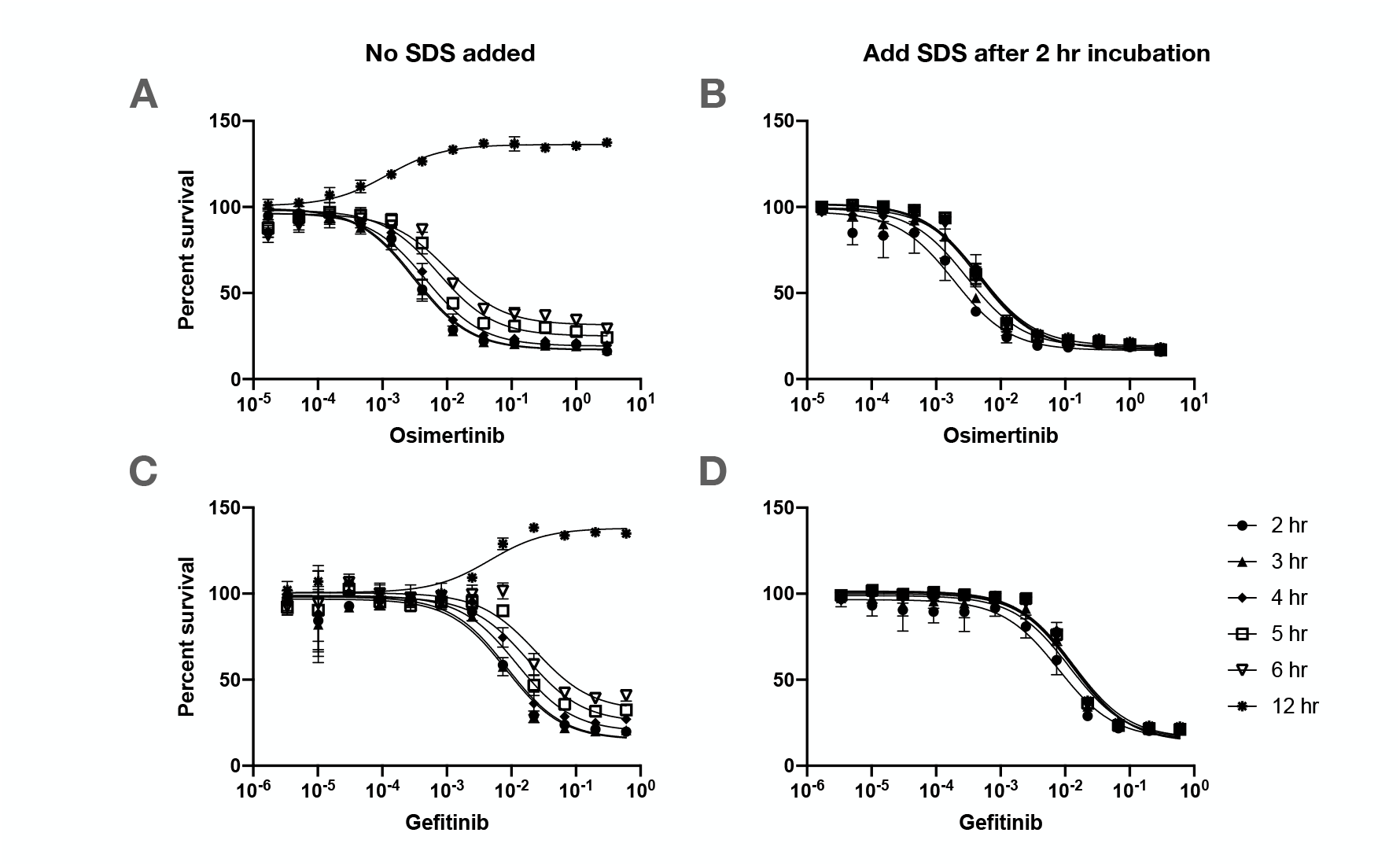
Adding SDS quenches the AB reaction and provides a way of preventing plate-to-plate variation in large scale experiments where multiple plates are to be read. PC-9 cells were treated with both varying concentrations of osimertinib (a & b) and gefitinib (c & d). After the 5 day drug incubation, AB was added and after the 2 hour conversion of AB, SDS was added at a final concentration of 1%. Readings at 2, 3, 4, 5, 6, and 8 hours show a right shift over time in the non SDS conditions. Mean + SD, n=3. **(a)** Osimertinib drug assay with Alamar Blue without SDS **(b)** Osimertinib drug assay with Alamar Blue with SDS **(c)** Gefitinib drug assay with Alamar Blue without SDS **(d)** Gefitinib drug assay with Alamar Blue with SDS.

### Calibration of fluorometer enables inter-plate comparison

Placing a common fluorescence standard solution into each plate for fluorometer calibration would ensure consistent readings and enables reliable plate-to-plate comparison even the comparison among the results obtained at different times. This capability is particularly important for determining the temporal change in drug sensitivity to understand drug resistance evolution upon exposure to therapeutics^8^. We first tested a pooled biologically converted resorufin as a calibration standard. To create this, we plated a high density of PC-9 cells (1 × 10^6^) in a 10 cm dish and added AB and let sit overnight. We then collected the media containing the biologically converted AB and used this as control wells in further drug assays. While the biologically converted resorufin is a good calibration standard, there may be batch to batch variations. In order to establish more consistent and convenient calibration standard, we tested Rhodamine B, a xanthene fluorescent dye, which has a similar excitation and emission spectrum profiles (ex/em 553/627 nm) as resorufin (ex/em 560/590 nm). We first made a stock of Rhodamine B solution in DMSO (1.8 mg/mL) and kept it at 4°C protected from light. On the day of measurement, we dilute the Rhodamine B stock solution with phosphate buffered saline (PBS) at 1:100 ratio by volume and the fluorescent intensity of 100 µl/well was determined by BioTek Synergy H1 (BioTek, Winoosi, VT) plate reader at 560/590nm excitation/emission with sensitivity setting at a fixed sensitivity setting at 72. Creating stock solutions every assay may be time consuming and a source of variability, so we tested and confirmed that the signal is stable at each measurement over time by diluting from a single source of stock solution. **Supplementary Figure S3** shows tested the fluorescence intensity up to 18 days, which confirms that the signal is stable over time and can be stored at 4°C protected from light.

Previous studies have also used Alamar Blue as a continuous monitoring assay. Kwack et al. showed Alamar Blue to be a non-radioactive method for measuring the bioactivity of interleukin (IL-2)^9^. They proved that Alamar Blue was comparable to that of the ^[3H]^thymidine incorporation method, a widely used protocol to monitor DNA synthesis and cell proliferation. Zhou et al. used Alamar Blue to track 3D bone tissue engineering constructs, showing that AB can be successfully used to monitor and predict cell confluence^10^. When non-destructive AB assays are repeated on the same plate with plate reader calibration as described in this paper, temporal cell viability changes of individual wells can be monitored longitudinally. We applied this longitudinal assay to an osimertinib drug sensitivity assay plate, and observed drug concentration dependent differential recovery kinetics (**Figure 4A & B**). We validated this method to be in correspondence CTG by measuring the cell proliferation kinetics, and this is shown in **Supplementary Figure S4**. Thus, AB has potential to be a good tool for continuous monitoring of cell viability changes.

**Figure 4.**
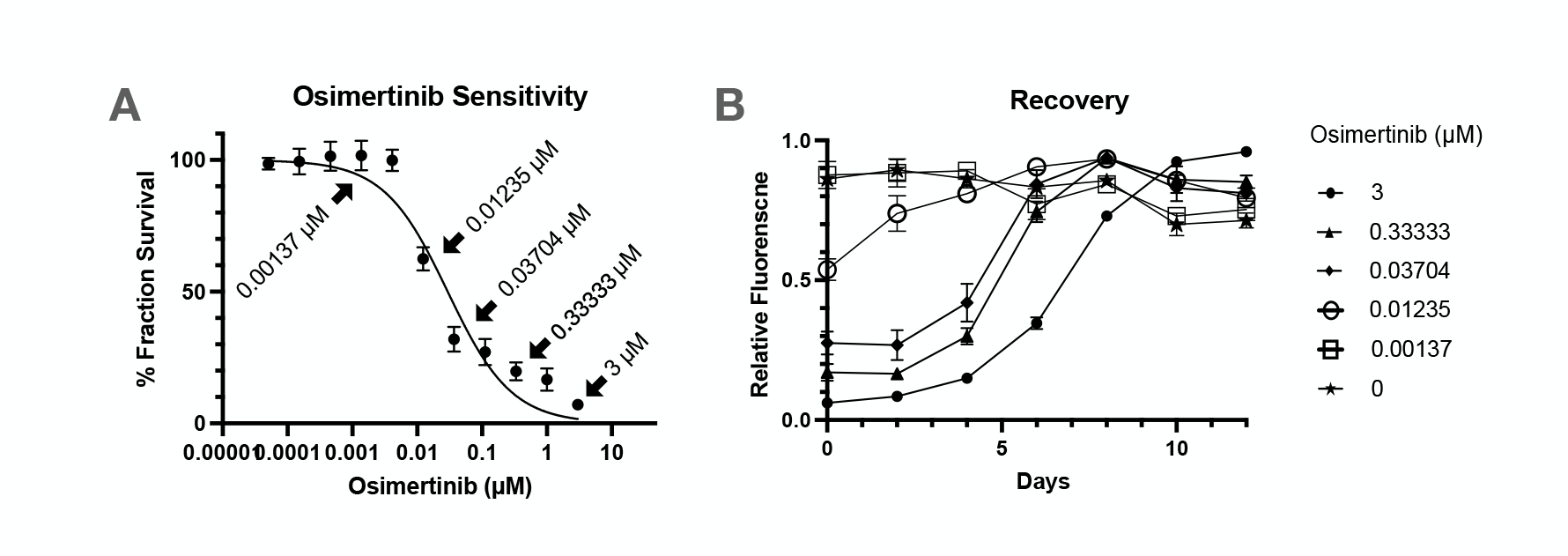
AB can be utilized as a longitudinal assay to monitor recovery following a drug sensitivity assay. **(a)** Drug sensitivity was determined by 2 hour Alamar Blue incubation following drug solution removal without SDS addition (n=6, mean + SD). The data points indicated by the arrow with drug concentrations are the samples illustrated in panel B for their recovery. **b** After fluorescence intensities were read, Alamar Blue reaction solution was washed off and cells were continued to cultured with fresh medium. Alamar Blue assay was repeated every two days.

## Discussion

We reported here that EGFR inhibitors can alter the florescence readings in AB based cell viability assays and create false readouts. We demonstrated that simple modifications to the published protocol may overcome these barriers and allow AB to be a feasible cost-effective alternative to other cell viability assays, such as CTG. Our protocol suggestions are as follows; 1) removing the media containing drug and add fresh media + 10% AB; 2) adding SDS to quench the reaction after a 2 hour incubation; and 3) utilizing a calibration standard to minimize plate-to-plate variation.

## Methods

### Materials

EGFRmut NSCLC PC-9 cells (Cat# 90071810-1VL), and Rhodamine B (Cat# 83689) were obtained from Millipore Sigma (St. Louis, MO). All 4 EGFR inhibitors: osimertinib (AZD 9291) (Cat# 16237), gefitinib (Cat# 13166), erlotinib (Cat# 10483, and lapatinib (Cat# 11493) were obtained CaymanChemical (Ann Arbor, MI). Alamar Blue(Cat# BUF012A) was purchased from Bio-Rad (Hercles, CA). Sodium dodecyl sulfate (Cat# 28312) was ordered from ThermoFisher (Waltham, MA). Cell-Titer Glo (Cat# G7572) was ordered from Promega (Madison, WI). Dimethyl sulfoxide (Cat# 276855) was ordered from Sigma-Aldrich (St. Louis, MO).

### Cell culture

PC-9 cells were maintained in Roswell Park Memorial Institute (RPMI) medium supplemented with 10% heat inactivated fetal bovine serum (FBS) and 1% streptomycin/penicillin in an incubator at 37°C under humidified 5% CO_2_ containing atmosphere.

### Drug Treatment

To set up drug assay plates, PC-9 cells were trypsinized from a 70-80% confluent stock culture, counted, and plated into 96-well plates at the density of 3000 cells/well in 90 *µ*L of medium and allowed to attach overnight. On the next day, drugs dissolved in DMSO were diluted by 3 fold serial dilution maintaining DMSO concentration constant (1% by volume) in culture medium. 10*µ*L of each resulting drug solution of a concentration of dilution series was transferred to each corresponding well of the drug assay plates seeded with PC-9 cells the day before. The final concentration of DMSO for treatment was 0.1% (v/v). After the five-day incubation, the cell viability was determined by Alamar Blue or CTG assay.

### Drug assay and analysis

For Alamar Blue assay with drug removal, the media containing drug was removed (discarded) and replaced with 100 *µ*L RPMI containing 10% (v/v). Control wells with no drug and background wells with no cells were also incubated with RPMI and Alamar Blue mixture For Alamar Blue assay without drug removal, 10 *µ*L of Alamar Blue solution was directly added to each well. Following 2-hour incubation in the CO_2_ incubator, 50 *µ*L of 3% (w/v) SDS in water was added to each well immediately to quench the reaction. Fluorescent signal was read by BioTek Synergy H1 (BioTek, Winoosi, VT) plate reader at 560/590nm excitation/emission.

CTG assay was performed according to the package instructions from Promega: 100µL of CTG was added to each well for a 1:1 CTG dilution in RPMI. After a 10 minute incubation time, luminescence was determined by BioTex Synergy H1 (BioTek, Winoosi, VT).

Data was analyzed using Prism GraphPad software (version 8.4.0 for macOS, GraphPad Software, San Diego, California USA, www.graphpad.com). The average background reading value determined from the readings of the wells without cells was subtracted from each reading to determined net signal. Each net reading value was normalized by average net value obtained from the wells without drug to calculate survival proportion. Curve fitting was done in Prism with dose-response inhibition, [inhibitor] vs. response (three parameters). Each drug concentration was tested in triplicate. Unpaired t-test was performed to compare IC50 values determined by multiplicated drug assays.

The survival fraction was calculated as follows:

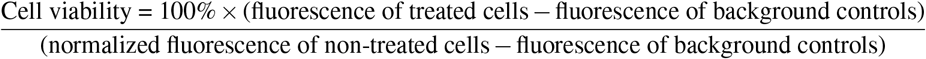

All 11 drugs and their highest concentrations tested are listed here. To find the IC50, we exposed the PC-9 cells to a 12 concentrations: 3 fold dilution starting from the highest concentration.

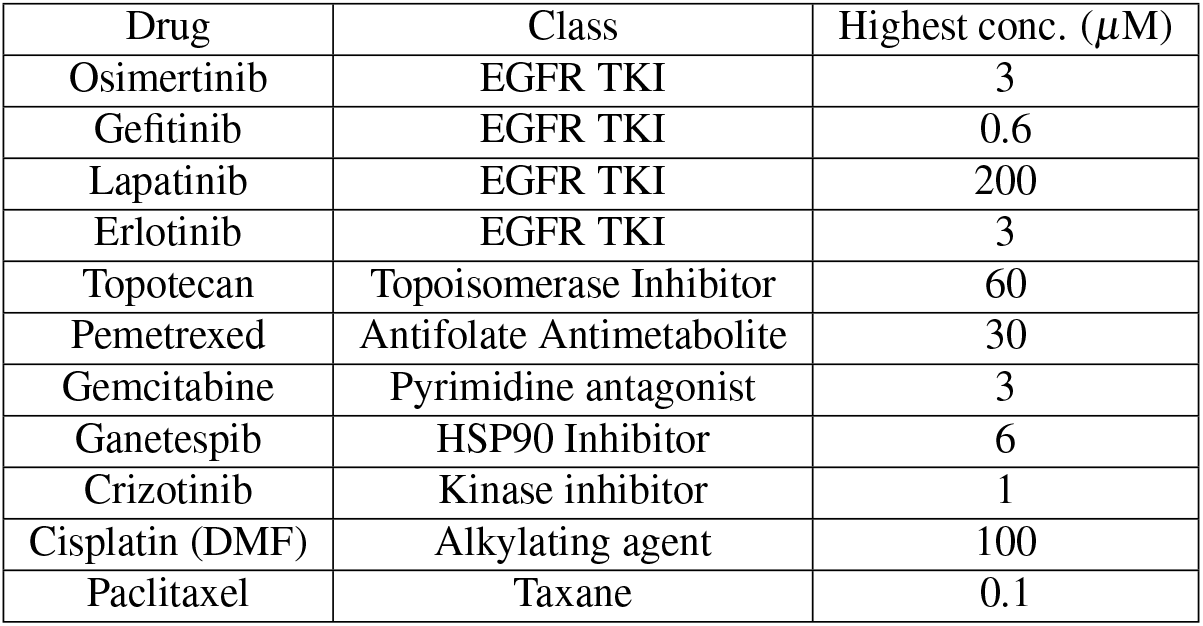

### Rhodamine B preparation

Rhodamine B calibration stock solution was prepared by diluting stock solution (1.6 mg / mL Rhodamine B in DMSO solution) was diluted with PBS by 100 fold prior to fluorometry. 100 µl of resulting solution was placed in wells of 96 well plates and the fluorescence intensity was measured using Synergy H1 hybrid plate reader with excitation at 560 nm, emission at 590 nm with sensitivity setting at 72.

## Supporting information

All supplemental figures

## Acknowledgements (not compulsory)

This study was supported by the National Institutes of Health (5R37CA244613-02) and the American Cancer Society Research Scholar Grant (RSG-20-096-01).

## Author contributions statement Additional information

To include, in this order: **Accession codes** (where applicable); **Competing interests** (mandatory statement).

The corresponding author is responsible for submitting a competing interests statement on behalf of all authors of the paper. This statement must be included in the submitted article file.

## Notes

### Competing Interest Statement

The authors have declared no competing interest.

