## Supplementary material for "Alamar Blue assay optimization to minimize drug interference and inter-assay viability": All supplemental figures

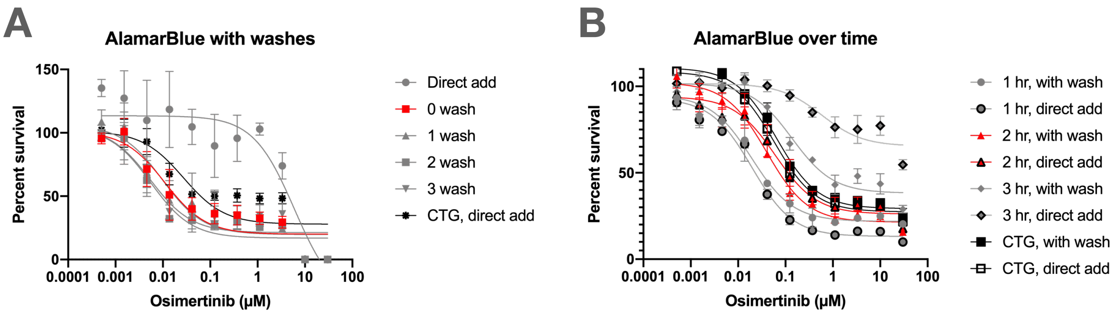


**Figure S1.** **Alamar Blue with no wash and a two hour incubation time is most similar to CTG.**
(**a**) There is no difference in the amount of washes. A wash is defined as dumping the media containing drug and replacing with 10% AB in fresh media. (**b**) Alamar Blue with no wash at an incubation time of 2 hours is most similar to CTG conditions.

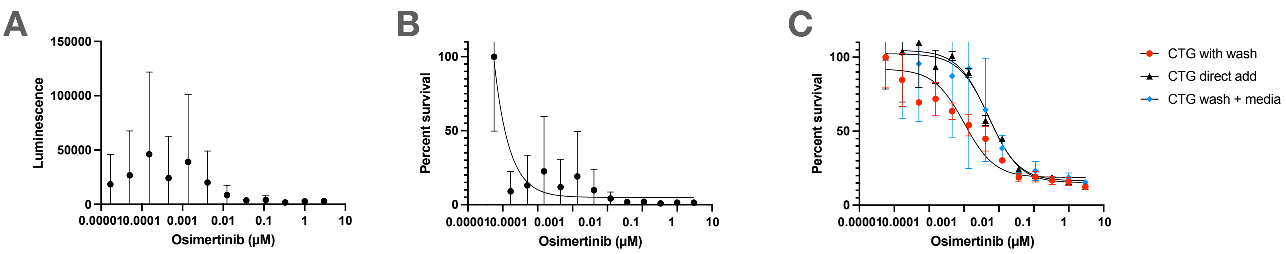


**Figure S2.** **CTG assay with and without wash show an expected drug curve shift.**
We tested CTG with wash and without wash to see if there was also an interference between osimertinib and the cell viability assay. We saw a similar left shift as when we used AB, but we hypothesized that this could be due to the exogenous ATP floating in the media that would be taken away during the ”dump” stage, since CTG utilizes ATP to quantify metabolically active cells. To test this, we transferred the media removed in the wash stage into a new 96 well plates and read the luminescence. Panel (**a**) shows the raw luminescence values and panel (**b**) shows the normalized values when divided by the no drug values (percent survival). (**c**) When we add the values of (a) and (b), the curve is equivalent to the CTG direct add condition.

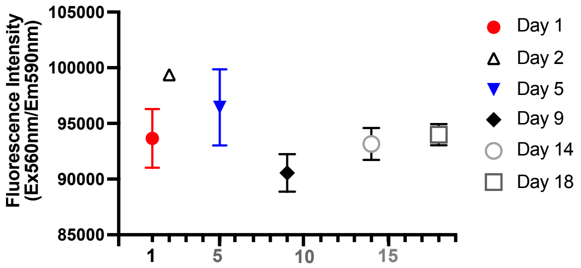


**Figure S3. Rhodamine B serves as a standard to calibrate fluorometer sensitivity.**
Freshly prepared Rhodamine B solutions with 3 independent dilutions on various days emitted consistent fluorescence intensities with minimal day-to-day variability, indicating it suitability to serve as a calibrator to set the plate reader sensitivity and as a reference standard for interplate comparison. Mean + SD, n=3.

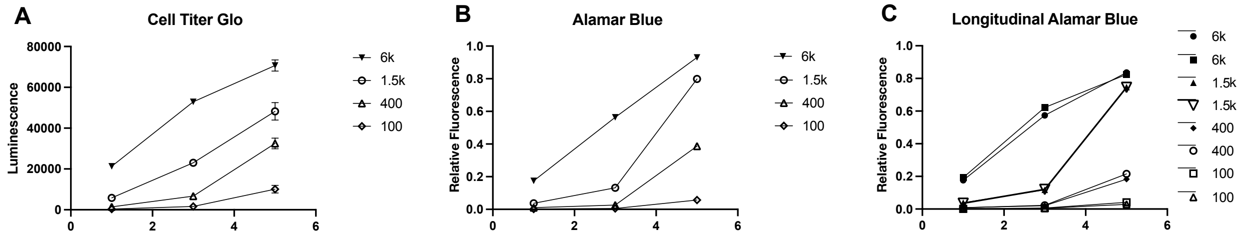


**Figure S4. PC9 cells growth curves determined by CTG, AB and longitudinal follow-up with AB.**
PC9 cells were plated into 96 well plates at indicted cell numbers per well. (**a** and **b**) Three sets of triplicated samples for each cell number were set to determine cell viabilities on day 1, 3, and 5 for CTG and AB assay (n=3, mean + S.D.). (**c**) Longitudinal cell proliferation kinetics of the culture in single wells were monitored by repeated Alamar Blue assays. The fluorescence intensity of each well was determined immediately after 2 hour-incubation with Alamar Blue. After reading Alamar Blue solution was washed off and cells were cultured in fresh medium until the next viability assay with Alamar Blue. The proliferation kinetics of PC9 cells with various initial seeding densities determined in this longitudinal fashion recapitulate that pattern determined by Alarmar Blue assay using triplicated samples for each time point (B).
